# Exploring *ex vivo* modulation of fibrosis in discarded human donor kidneys

**DOI:** 10.1101/2025.07.31.667749

**Authors:** C.C. Pamplona, C.B. Bergada, C.L. Jaynes, T.L. Hamelink, V.A. Lantinga, B. Ogurlu, P Ottens, K. van Hateren, J.L. Hillebrands, P. Olinga, D Touw, H.G.D. Leuvenink, C. Moers, L.L. van Leeuwen

## Abstract

**Introduction:** Early-onset fibrosis limits kidney transplant success. Normothermic machine perfusion (NMP) offers a platform for targeted drug delivery directly to isolated organs, minimizing systemic effects. This study evaluated the long-term anti-fibrotic efficacy and safety of galunisertib in discarded human kidneys perfused *ex vivo*.

**Methods:** Twelve discarded human kidneys underwent 4 hours of oxygenated hypothermic perfusion followed by 6 hours of NMP with galunisertib or vehicle (n=6). Precision-cut kidney slices (PCKS) were then cultured for 48 hours with either continued or discontinued galunisertib exposure. Endpoints included fibrosis-related mRNA expression and pharmacokinetics.

**Results:** Galunisertib did not negatively affect renal function during NMP. Continued exposure in PCKS significantly attenuated fibrosis-related mRNA expression, including SERPINE1 (p=0.0046), TGF-β (p=0.0168), FN1 (p=0.0269) and ACTA2 (p=0.0014) after 48 hours. The discontinuation of treatment did not exhibit the same anti-fibrotic effects.

**Conclusion:** Galunisertib was safely administered during NMP and steadily excreted via the urine. NMP showed to be a promising platform for safe targeted anti-fibrotic therapy delivery, offering potential to improve graft quality. When treatment was sustained, galunisertib induced a modest reduction in fibrosis-related mRNA expression over 48 hours of tissue incubation. Further studies are needed to optimize delivery strategies and evaluate the impact of prolonged therapeutic exposure.

## Introduction

Chronic allograft nephropathy, a progressive condition marked by interstitial fibrosis and tubular atrophy, is the most common cause of graft failure. In average, only 50% of kidneys from deceased donors remain functional 10 years post-transplantation, with early-onset fibrosis being the leading cause of graft failure.^1,2^ During ischemia-reperfusion injury (IRI), a cascade of damage in kidney grafts is initiated, triggering cell death, inflammation, and structural alterations that compromise tissue integrity. IRI disrupts endothelial cells, depletes the glycocalyx, and destabilizes the cytoskeleton, leading to vascular and tubular injury. In response, the kidney activates wound-healing processes aimed at restoring function; However, under prolonged or dysregulated conditions, this response shifts toward pathological fibrogenesis. This phenomenon involves the gradual replacement of functional renal units with scar tissue, leading to a continuous decline in kidney function. It often starts as a result of IRI or pre-existing fibrosis in the donor kidney and is exacerbated by the alloimmune response, therefore amplifying inflammation, endothelial dysfunction and, ultimately, contributing to compromised graft quality before transplantation. ^1–3^

The upregulation of transforming growth factor-beta (TGF-β) upon ischemic injury plays a central role in fibrogenesis by activating the small mother against decapentaplegic (SMAD) signaling pathway. Chronic inflammation and excessive activation of this cytokine lead to an overproduction of extracellular matrix proteins in the tissue, progressively altering the organ’s structure and function. ^1,4,5^ Ultimately, fibrosis emerges as the final, irreversible stage of injury to the vasculature and renal tissue.

*Ex vivo* normothermic machine perfusion (NMP) has become a commonly used technique for preservation, resuscitation, and functionality assessment of suboptimal donor organs prior to transplantation. ^6–11^ The implementation of machine perfusion after warm ischemic injury is currently the most promising technique of counteracting the early onset of fibrosis, not only because it improves organ preservation but also its potential for pioneering targeted drug delivery to an isolated organ, thereby eliminating the risk of systemic adverse effects. ^2,3,12^ Currently, only a limited number of therapies are available that may attenuate fibrogenesis, yet their feasibility and effectiveness in human kidneys remain to be determined.

Galunisertib is a selective inhibitor of the SMAD signaling pathway that targets the TGF-β type I/II receptors. Originally developed as an anti-neoplastic agent, it has since demonstrated potent anti-fibrotic effects in multiple studies through its inhibition of TGF-β signaling.^1,13^ When combined with NMP, galunisertib offers the potential for targeted inhibition of fibrosis within kidney grafts, while minimizing systemic side effects and potentially enabling the use of lower therapeutic doses.

In prior studies using a porcine model, our team demonstrated that repurposing galunisertib as an anti-fibrotic agent during NMP led to pronounced and sustained anti-fibrotic effects, which remained evident for 48 hours in subsequent tissue slice analyses. ^2,3,14^ Building on these compelling findings, we sought to translate this approach to a preclinical setting by administering galunisertib to discarded human kidneys during *ex vivo* NMP, followed by tissue slice incubation to evaluate its long-term effects on fibrinogenesis. With this approach, we aim to eventually enhance graft quality and extend graft survival by mitigating the development of fibrosis. ^15,16^

## Methods

### Study Design

This study aimed to investigate the anti-fibrotic effect of galunisertib on donor kidneys using a combination of normothermic machine perfusion and precision-cut kidney slices (PCKS).

Twelve human discarded kidneys were randomly assigned to two groups, namely a control group and a galunisertib-treated group. Upon arrival at the laboratory on static cold storage (SCS), all kidneys underwent oxygenated hypothermic machine perfusion (HMP-O_2_) for 4h followed by 6 hours of NMP. In the galunisertib treated group, the drug was added to the reservoir 1 hour after the start of NMP, while the control kidneys received only the vehicle.

After NMP, kidneys were flushed with cold preservation solution and 6mm core biopsies were prepared from the cortex for future tissue slicing. The PCKS were then preserved in an oxygenated incubator for 48 hours undergoing continuous gentle agitation. The galunisertib-treated kidneys were further divided into two subgroups: one in which the treatment was continued throughout the whole incubation period and another in which treatment was discontinued. The control kidneys remained untreated for the entire duration of incubation.

Our primary outcome measure was the effect of galunisertib on gene and protein expression of fibrosis-related markers. Secondary endpoints were the toxicity and safety of galunisertib on renal function, vitality, and injury. These parameters were assessed via common functional NMP parameters as well as oxygen consumption, ATP levels in the renal cortex biopsies and tissue slices, and injury markers during NMP and histological staining throughout the experiment. Investigators were blinded for the experimental group up until the organs were in HMP, but not blinded during the execution of normothermic machine perfusion and slice experiments as solutions needed to be specifically made. Nonetheless, the analysis was performed blinded.

### Donor cohort

Deceased donor kidneys deemed unsuitable for transplantation were included in this study. These kidneys exhausted United Network for Organ Sharing (UNOS) allocation after no suitable recipients were identified due to poor quality and/or logistical issues. Research use was approved by next-of-kin consent, and cold ischemia time was maintained below 40 hours as a target criterion. This study aimed to include deceased donors with higher Kidney Donor Profile Index (KDPI), glomeruli sclerosis, fibrosis scores, and arterial sclerosis during donor/kidney pathological assessment.

Exclusion criteria included organs with aberrant arterial anatomy during organ retrieval that impeded cannulation for perfusion, donor virology positive for Hepatitis A/B/C and HIV, a cold ischemia time at arrival above 40 hours, technical issues during NMP that resulted in >10 minutes of warm ischemia time (WIT), macroscopically visible ischemia areas during NMP, and infections during slice incubation. These criteria were established prospectively.

### Organ Procurement and Preservation

Organ retrieval was performed according to US standards, and kidneys were then shipped to the laboratory. Upon arrival, the kidneys were (re)cannulated and connected to a custom-built HMP device and a Kidney Assist Transport® (XVIVO, Göteborg, Sweden) disposable. HMP-O_2_ was performed with 330mL University of Wisconsin Machine Perfusion Solution (UW-MP) for 4 hours. The setup consisted of temperature control between 0-4°C, a pulsatile pressure-controlled perfusion setup with a mean arterial pressure of around 25□mmHg, and 100% oxygenation at a flow rate of 0.1mL/min.

### Normothermic Machine perfusion

After completion of HMP, kidneys were connected to an open-circuit NMP setup (without venous cannulation) as previously described by our group. ^8,17^ The perfusate composition is described in Table 1. The circuit was primed with an oxygenated acellular perfusion solution (Table 1). Red blood cells were added in when the temperature reached approximately 30°C, to avoid cellular injury. Oxygenation was achieved using carbogen (95% O_2_ / 5% CO_2_) at a flow rate of 0.5 L/min.

**Table 1:** Composition of solutions throughout the experiment.

| COMPONENT | AMOUNT |
| --- | --- |
| **Perfusate** |  |
| *Washed O-neg RBCs with NaCl 0.9% (Fresenius Kabi)* | 418 mL |
| *Cernevit multivitamins (Baxter B.V., Utrecht, The Netherlands) dissolved in 5mL NaCl 0.9%* | 0.5 mL |
| *Cefazolin 1g (Sandoz B.V., Almere, The Netherlands) dissolved in 10 ml sterile water* | 10 mL |
| *Heparin 25000IU/mL (Sagent Pharmaceuticals, Schaumburg IL, USA)* | 1250 IU |
| *Calcium gluconate 10% (B. Braun, Melsungen AG, Melsungen, Germany)* | 4 mL |
| *Sodium bicarbonate 8.4% (B. Braun, Melsungen AG, Melsungen, Germany )* | 20 mL |
| *Glucose 25% (Pfizer Inc., New York, USA)* | 2.8 mL |
| *Mannitol 25% (Fresenius Kabi)* | 3 mL |
| *Magnesium sulfate 100 mg/mL (Teva Nederland B.V., Haarlem, The Netherlands)* | 0.5 mL |
| *Aminoplasma 10% (B. Braun)* | 5 mL |
| *Sodium phosphate 3 mmol/mL (Apotheek A15, Gorichem, The Netherlands)* | 0.15 mL |
| *Human albumin 20g/L (Sanquin Plasma Products B.V., Amsterdam, The Netherlands)* | 100 mL |
| *Sterile water (Fresenius Kabi)* | 113.3 mL |
| *Creatinine (Sigma-Aldrich, Zwijndrecht, The Netherlands)* | 0.052 gr |
| **Infusion Solutions** |  |
| *Flolan 0.5mg (GlaxoSmithKline, Middlesex, United Kingdom) dissolved in NaCl 0.9% solution* | 4 μg/h |
| Glucose 5% (diluted from 25% in sterile water) | 3 mL/h (if necessary) |
| **Culture Medium (per 100mL)** |  |
| Sterile Williams E medium (Glutamax) (ThermoFisher Scientific)<br>*diluted from Sterile Williams E medium powder resuspended in 973.81 mL sterile water + 26.19 mL of sodium bicarbonate 8.4% solution | 100mL |
| Ciproflaxine (Sigma-Aldrich) | 6.03μM |
| Fungizone (Sigma-Aldrich) | 0.027nM |
| Glucose powder 180.16 g/mol (Sigma-Aldrich) | 1.52mM |
| **1x Krebs Solution (per 1L)** |  |
| 10x concentrated Krebs solution in milliQ water<br>* Calcium chloride powder (CaCl<sub>2</sub>) – 25mM (Sigma-Aldrich)<br>Potassium chloride powder (KCl) – 50mM (Sigma-Aldrich)<br>Sodium chloride powder (NaCl) – 1.18M (Sigma-Aldrich)<br>Magnesium sulfate powder (MgSO) – 11mM (Sigma-Aldrich)<br>Potassium dihydrogen phosphate powder (KH) – 12mM (Sigma-Aldrich) | 100 mL |
| NaHCO<sub>3</sub> powder (Sigma-Aldrich) | 25mM |
| Glucose powder (Sigma-Aldrich) | 25mM |
| HEPES powder (Sigma-Aldrich) | 10mM |
| Cold Mili-Q (Sigma-Aldrich) | 900 mL |

From the start of perfusion, a continuous infusion solution of epoprostenol (Flolan 0.5mg at 4 µg/h) was administered (Table 1). Kidneys were perfused for 6 hours with a sinusoidal pressure of 110/70mmHg and temperature of 37°C. Arterial pH and glucose concentration were adjusted prior to start of NMP and hourly thereafter based on the measured blood gas values. If pH dropped below 7.3, sodium bicarbonate (8.4%) was administered to maintain a target range of 7.35–7.45. Glucose levels were corrected with 5% glucose solution to maintain a final concentration of 6 mmol/L when levels fell below 4 mmol/L. In case of rapid glucose depletion to below 5mmol/L within the first 2 hours, an infusion solution of 5% glucose was initiated at 3mL/h until stabilization (Table 1). The renal arterial pressure, flow rate, temperature, and urine production were continuously logged throughout NMP. At the 60-minute time point, galunisertib (2.6 mM Axon Medchem, LY2157299) (Sigma-Aldrich, Zwijndrecht, The Netherlands) was added to the reservoir diluted in 2.61 mL DMSO for a final concentration of 10μM. For control kidneys, the same volume of DMSO (2.61 mL) was administered, obtaining a perfusate DMSO concentration of 0.38%.

### Precision-cut Kidney Slices (PCKS)

PCKS were prepared as previously described by our group. ^3^ In short, the kidneys were immediately flushed with 300mL of cold (0-4°C) University of Wisconsin cold storage (UW-CS) solution after ending NMP. Subsequently, cortical tissue biopsies were made using a 6mm punch biopsy. The tissue cores were then transferred to cold UW-CS solution. With a Krumdiek Slicer (Alabama Research and Development, Munford, AL, USA) cooled to 4°C with a water bath and primed with oxygenated 1x concentrated Krebs Solution (pH 7.4), cortical renal tissue slices were cut with a thickness of around 300□μm.

Slices were cultured in pre-warmed (37°C) and oxygenated (5% CO_2_ and 80% O_2_) 12-well plates containing culture medium (1.3□mL per well) for 48 hours while being gently shaken at 90□cycles/min, with a medium refresh at the 24-hours timepoint. Culture medium components are described in Table 1. To assess whether the effects of galunisertib persisted beyond NMP, the six treated kidneys were subsequently divided into two groups during the PCKS phase: one with continued galunisertib treatment and one with treatment discontinued (refer to study design, Figure 1). For the continued treatment group, 10µM galunisertib was added to the medium diluted in DMSO (5µL), while the discontinued treatment group was supplemented with 5µL of DMSO.

**Figure 1:**
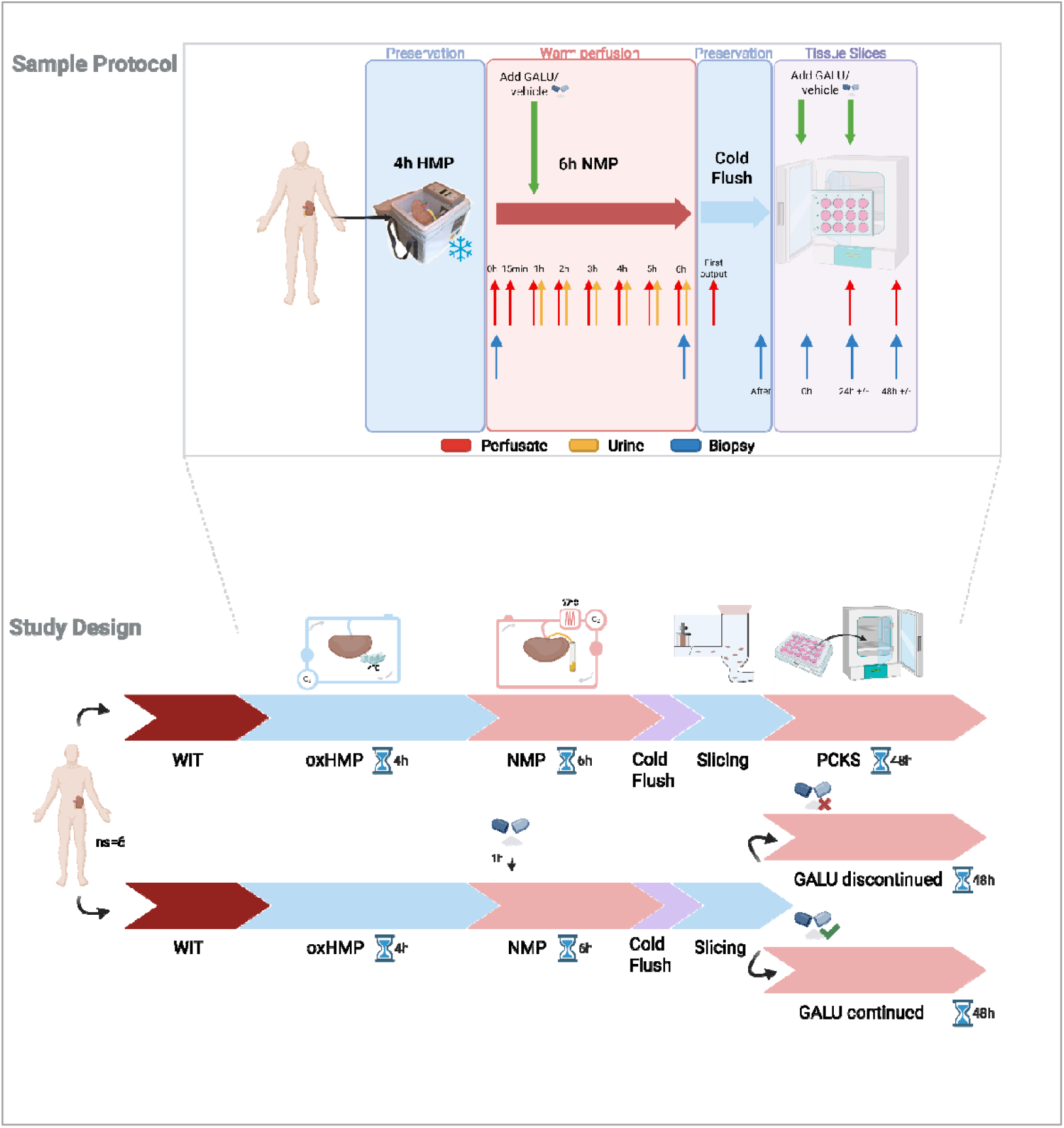
Study Design and Sample Protocol. Twelve discarded human kidneys were randomly assigned to control/vehicle or galunisertib-treated groups. After arrival on static cold storage, all kidneys underwent 4 hours of oxygenated hypothermic machine perfusion (HMP-O_2_) followed by 6 hours of normothermic machine perfusion (NMP). Galunisertib or vehicle was added after 1 hour of NMP. Post-NMP, kidneys were flushed, and 61lmm cortical biopsies were taken for precision-cut kidney slices (PCKS), which were incubated for 48⍰h at 37°C. During incubation, galunisertib treated kidneys were subsequently divided into subgroups with either continued or discontinued treatment.

### Sampling and Analysis

#### Perfusate, Urine, Flush, and Medium

Arterial and venous perfusate samples were collected for blood gas analysis at baseline (prior to NMP), at 30 and 60 minutes (before treatment), and then hourly throughout the remainder of the perfusion. Blood gas measurements included pH, partial oxygen pressure, oxygen saturation, hemoglobin, glucose, and were used to calculate oxygen consumption. Additional perfusate “plasma” samples were taken and stored for future quantification of cellular injury (aspartate aminotransferase (ASAT) and lactate dehydrogenase (LDH)), renal filtration (creatinine clearance (CrCl) and fractional sodium excretion (FENa)), pro-collagen 1a1, TNFα, and IL-6 quantification, and for pharmacokinetic analysis of galunisertib (methods described in Supplementary Material).

Urine samples were collected to quantify urine output, perform pharmacokinetic analysis of galunisertib, and calculate creatinine clearance (CrCl) and fractional excretion of sodium (FENa). Sampling timepoints included 30 and 60 minutes (prior to treatment), followed by hourly intervals until the end of perfusion. For each timepoint, urine was collected during the final 10 minutes of the hour, with the remaining volume recirculated back into the reservoir. This approach preserved perfusate stability throughout most of the perfusion period while ensuring urine samples accurately reflected recent physiological conditions, synchronized with concurrently collected blood samples.

A flush sample was taken from the veins at the start of the post-NMP flush, allowing the measurement of galunisertib washed out of the kidney after perfusion.

During tissue slice incubation, medium samples were taken at 24h and 48h for pharmacokinetic analysis and pro-collagen 1a1 quantification. For an overview of all collected samples, refer to Figure 1 – Sample Protocol.

#### Biopsies

Biopsies and PCKS were collected for quantification of ATP and fibrosis gene/protein expression (methods described in Supplementary Material). Fibrosis-related expression included quantification of *TGF-β, SERPINE1, SERPINH1, FN1*, and *ACTA2*. These genes were selected due their short turnover and to confirm previous finding from our group in a similar porcine mode l. ^3^ For this analysis, a house-keeping gene was selected for fold-induction analysis, and its stability throughout the assay was considered and checked.

During NMP, cortical punch (4mm) biopsies were taken from the upper pole before connection to the circuit, at the end of warm perfusion, and after the post-NMP flush. PCKS samples were collected before incubation, at 24h, and at 48h. For analyses requiring tissue slices, three slices were pooled for each measurement.

### Statistics

The study was designed with equal-sized groups, assigned through randomization. Data visualization and analysis were performed using GraphPad Prism (Version 8.4.2). Values are shown as the mean + standard deviation (SD). For longitudinal data, variations among all experimental groups were evaluated using a two-way analysis of variance (ANOVA). Pairwise comparisons between dependent groups were conducted using a paired t-test or, when normality assumptions were not met, the Wilcoxon matched-pairs signed rank test. Comparisons between independent groups were performed using an unpaired t-test, or the Mann-Whitney test for non-normally distributed data. Donor characteristics between the groups were compared using table1_mc command in STATA (Version 18.5). Continuous variables were reported as means with standard deviation and compared using the t-test while categorical variables were presented as counts with percentages and compared using chi-square test of Fisher’s exact test. During PCKS analysis, control values were repeated at the galunisertib continued treatment groups at 24 hours and 48 hours for comparison, as the control NMP groups did not undergo treatment during PCKS. All statistical tests were two-sided, A *p*-value <□0.05 was assumed to indicate statistical significance. Flow, urine output, oxygen consumption, and CrCl data were normalized for kidney weight, ATP measurements were normalized per gram of protein, and gene expression was normalized for fold induction of house-keeping gene ACTB to minimize unwanted sources of variation.

## Results

### Donor Characteristics

Donor kidneys included in the study were from deceased donors aged 47–67 years, with a Kidney Donor Profile Index (KDPI) between 75–99. Histopathological assessments revealed glomerulosclerosis scores ranging from 6–73%, fibrosis scores between 10–25%, and arterial sclerosis between 11–50%. As shown in Table 3, there were no significant differences between the control and galunisertib-treated groups in terms of donor age, gender, kidney laterality, type of donation, last reported creatinine, warm and cold ischemia times (WIT/CIT), histological evaluations of fibrosis and sclerosis, or comorbidities such as hypertension and diabetes mellitus.

**Table 3:** Donor characteristics of the study groups.

| Variable | Control (N=6) | Galunisertib (N=6) | p-value |
| --- | --- | --- | --- |
| **Sex** |  |  | 0.22 |
| - Female | 3 (50%) | 1 (17%) |  |
| - Male | 3 (50%) | 5 (83%) |  |
| **Age (years)** | 58.8 (49-67) | 59.5 (47-67) | 0.87 |
| **Donation** |  |  | 0.08 |
| - DBD | 2 (33%) | 5 (83%) |  |
| - DCD | 4 (67%) | 1 (17%) |  |
| **KDPI score (%)** | 83.17 (54-95) | 90.17 (78-97) | 0.35 |
| **Last creatinine (mg/dL)** | 1.48 (0.49-2.3) | 2.61 (1-7.81) | 0.21 |
| **Laterality** |  |  | 0.56 |
| - Left | 3 (50%) | 4 (67%) |  |
| - Right | 3 (50%) | 2 (33%) |  |
| **Renal Resistance on HMP**<br>**(mmHg·min<sup>-1</sup>·mL<sup>-1</sup>)** | 0.33 (0.13-0.57) | 0.29 (0.12-0.53) | 0.69 |
| **Sclerosis (%)** | 14.67 (8-35) | 19.5 (2-46) | 0.55 |
| **Warm ischemic time (minutes)** | 20.98 (0-30) | 27.00 (0-27) | 0.43 |
| **Cold ischemic time (hours)** | 30.47 (21.47-42.53) | 23.95 (11.8-33) | 0.19 |
| **Hypertension** | 4 (67%) | 5 (83%) | 0.5 |
| **Diabetes Mellitus** | 1 (17%) | 3 (50%) | 0.22 |

### Functional perfusion parameters

Overall organ function following *ex vivo* administration of galunisertib was assessed. Parameters measured during HMP are shown in Figure S1. Functional perfusion parameters, including renal arterial flow, vascular resistance, urine output, arterial pH, creatinine clearance, and fractional sodium excretion during NMP (Figures 2A– G) showed no significant differences between the treatment and control groups.

**Figure 2:**
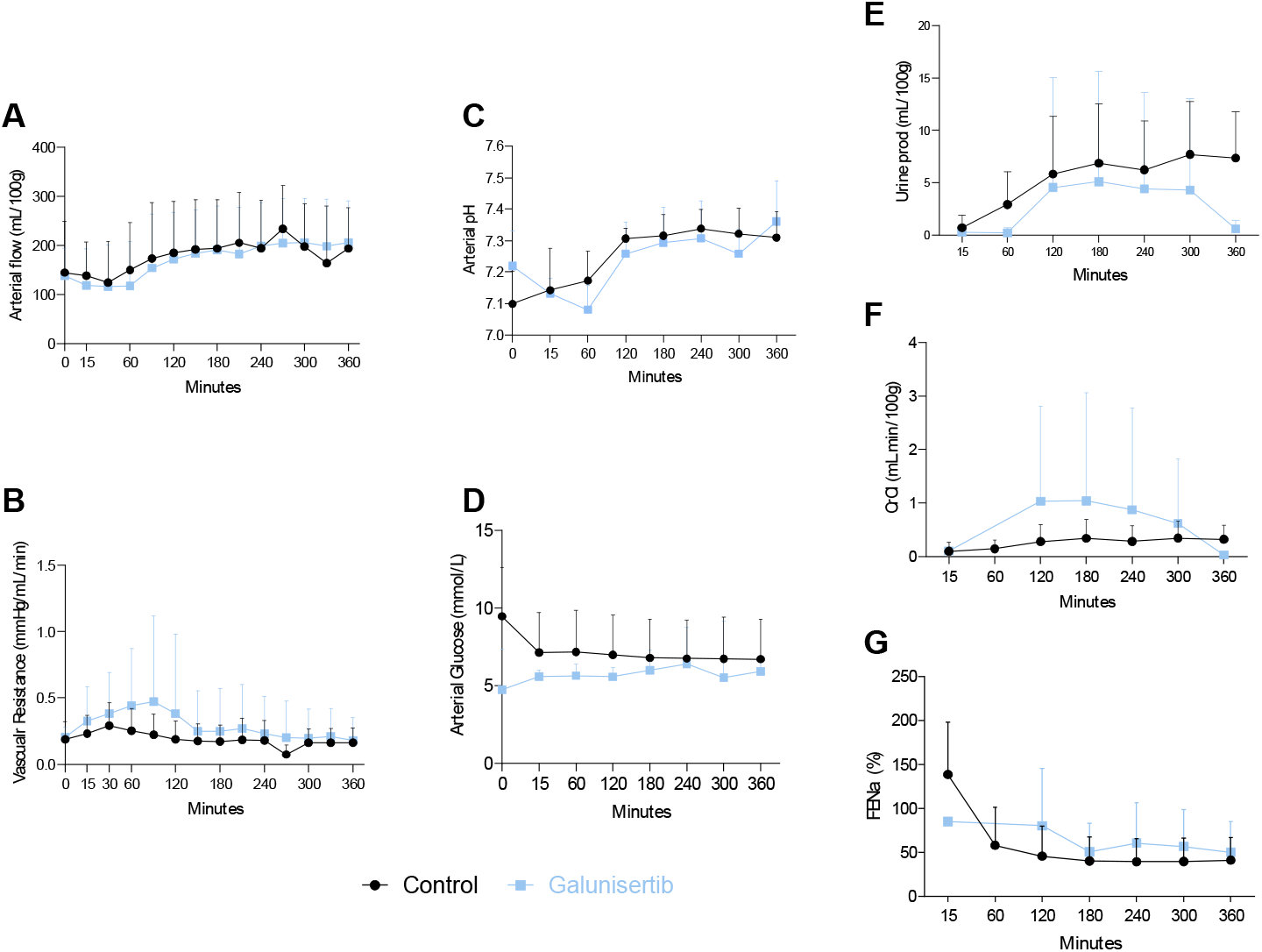
Overview of functional markers during 360 minutes of normothermic machine perfusion (NMP). (A) Grouped longitudinal representation of renal arterial flow (mL/min) per 100g of tissue; (B) Grouped longitudinal representation of vascular resistance (mmHg/mL/min); (C) Grouped longitudinal representation of arterial pH; (D) Grouped longitudinal representation of arterial glucose (mmol/L); (E) Grouped longitudinal representation of urine output (mL) per 100g of tissue; (F) Grouped representation of calculated creatinine clearance (CrCl) (mL/min) per 100g of tissue; (G) Grouped longitudinal representation of calculated fractional sodium excretion (FENa).

### Injury and Metabolic parameters

Subsequently, the effects on general injury markers and metabolism was investigated. As shown in Figures 3 A-B, biochemical injury markers remained similar throughout perfusion of both experimental groups, with a modest but non-significant increase over time. As for the metabolic parameters (Figure 3C and E), oxygen consumption was initially elevated during the warm-up phase and stabilized during the course of NMP in both groups. Renal tissue ATP levels before and after perfusion were similar across groups. Both groups showed an increase in TNFα and IL-6 expression over the 6-hour perfusion period; however, no significant differences were observed (Figure 3F-G).

**Figure 3:**
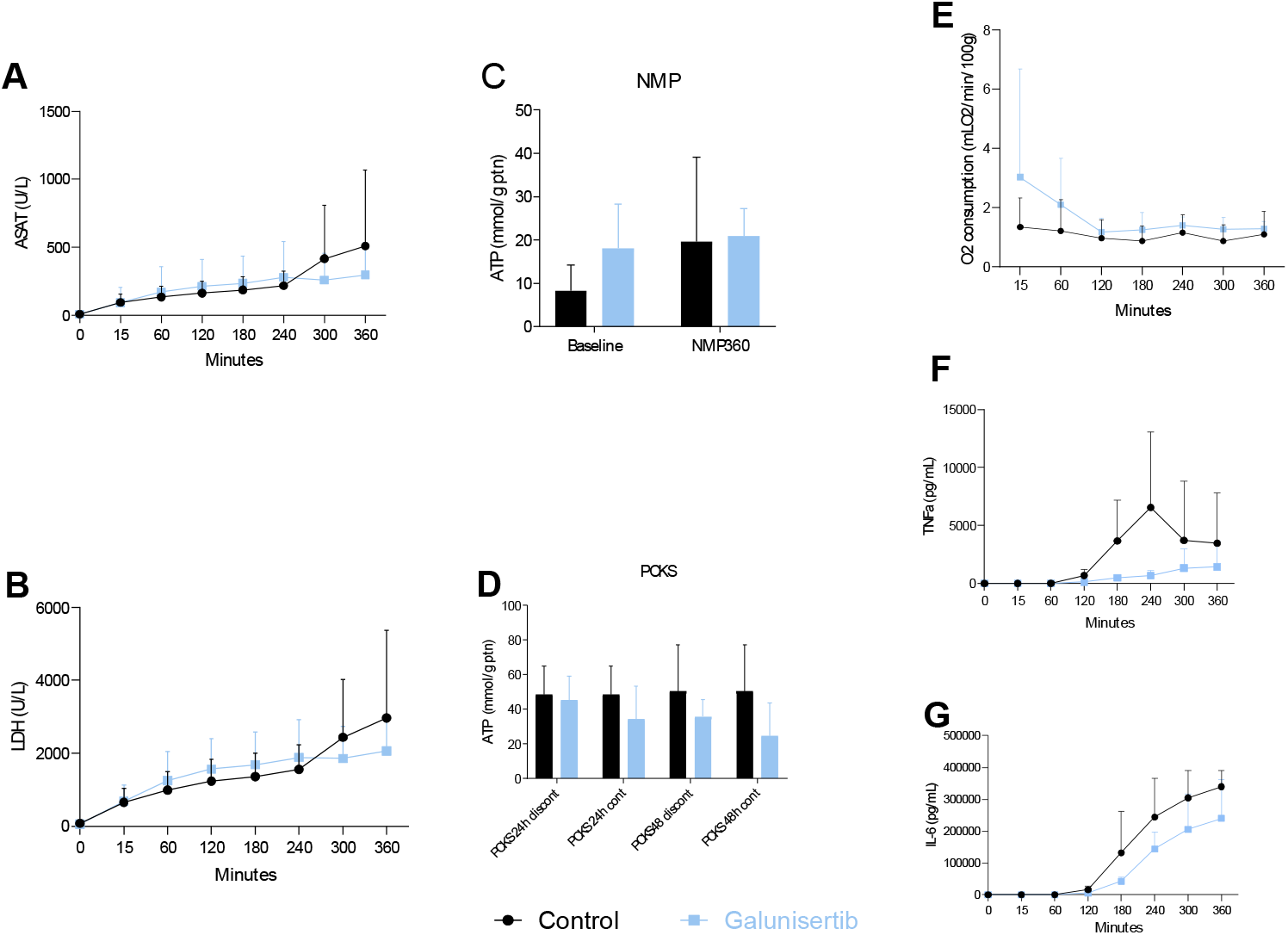
Overview of injury and metabolic markers during 360 minutes of normothermic machine perfusion and precision-cut kidney slices. (A) Grouped longitudinal representation of aspartate aminotransferase (ASAT (U/L); (B) Grouped longitudinal representation of lactate dehydrogenase (LDH) (U/L); (C) Representation of cortical ATP tissue levels at start and end of NMP (mmol/gram of protein); (D) Representation of PCKS ATP levels at 24 and 48 hours of both continued and discontinued treatment groups from kidneys perfused with galunisertib (mmol/gram of protein); (E) Grouped longitudinal representation of calculated oxygen consumption (VO2) (mLO_2_/min) per 100g of tissue; (F) Grouped longitudinal representation of TNFα in perfusate (pg/mL); (G) Grouped longitudinal representation of IL-6 in perfusate (pg/mL).

### PCKS Viability

ATP levels during PCKS incubation showed viability of slices throughout the whole experiment, with no significant differences between groups and timepoints.

### Fibrosis Gene expression and Protein secretion

For our primary outcome measure, we examined the short-term (during NMP) and long-term (during tissue slice incubation) effect of galunisertib on fibrogenesis. As shown in figure 4A, *TGF-β* expression did not differ between groups during NMP. However, sustained treatment during tissue slice incubation resulted in a significant reduction in *TGF-β* levels over time and compared to the control group (p=0.0168).

**Figure 4:**
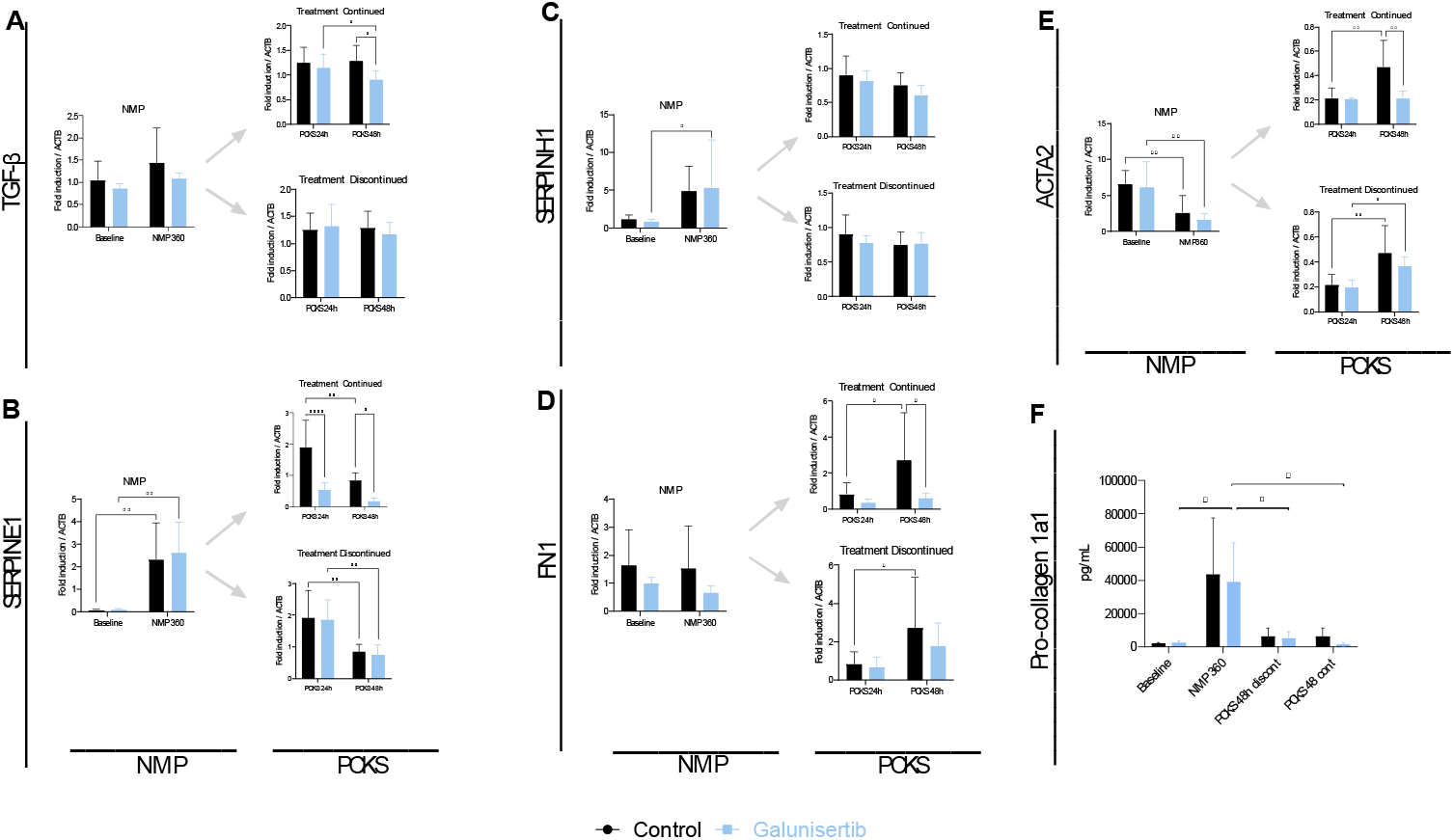
Overview of fibrosis-related mRNA in tissue and pro-collagen 1a1 protein secreted in perfusate during normothermic machine perfusion and precision-cut kidney slices (continued and discontinued treatment). (A) Representation of *TGF-β* mRNA expression levels in control and galunisertib treated kidneys represented in fold inductions from housekeeping gene *ACTB*; (B) Representation of *SERPINE1* mRNA expression levels in control and galunisertib treated kidneys represented in fold inductions from housekeeping gene *ACTB*; (C) Representation of *SERPINH1* mRNA expression levels in control and galunisertib treated kidneys represented in fold inductions from housekeeping gene *ACTB*; (D) Representation of *FN1* mRNA expression levels in control and galunisertib treated kidneys represented in fold inductions from housekeeping gene *ACTB*; (E) Representation of *ACTA2* mRNA expression levels in control and galunisertib treated kidneys represented in fold inductions from housekeeping gene *ACTB*; (F) Representation of pro-collagen 1a1 protein secreted levels in perfusate during NMP (baseline and 360 minutes of NMP) and medium at 48h of incubation from control and continued/discontinued galunisertib treatment (pg/mL).

TGF-β-associated transcripts, including *SERPINE1, SERPINH1, FN1*, and *ACTA2*, were quantified (Figures 4B-E). *SERPINE1* mRNA expression significantly increased during perfusion in both groups (*p*=0.0039 control and *p*=0.0018 galunisertib treated), with no significant differences observed between the treatment groups (Figure 4B). During tissue slice incubation, the expression significantly decreased in the control and discontinued treatment group, and continued galunisertib treatment led to a significant reduction in *SERPINE1* expression over time at both 24 and 48h (*p*=0.0057 at 24h and *p*=0.0046 at 48h).

In contrast, *SERPINH1* significantly increased during NMP in the galunisertib treated group (*p*=0.0464), while no differences were observed during tissue slice incubation. *FN1* levels did not significantly differ during perfusion, but an increase in the control group was observed during tissue slice incubation (*p*=0.0182). Furthermore, continued galunisertib treatment led to significantly lower *FN1* mRNA expression levels after 48h incubation compared to control (*p*=0.0269). Lastly, *ACTA2* mRNA expression reduced significantly during NMP in both control and galunisertib treated kidneys (*p*=0.0053 in the control group, and *p*=0.0026 in the treated group). Nevertheless, halting the treatment during PCKS caused significant increases in *ACTA2* expression in slices with discontinued treatment (*p*=0.0156), while continued exposure to the drug avoided this phenomenon and triggered a significant reduction in *ACTA2* expression between control and galunisertib treated slices (*p*=0.0014). To evaluate overall expression trends per kidney, we performed a longitudinal analysis of mRNA expression levels (Figure S2). Kidneys that received continuous galunisertib treatment throughout incubation exhibited generally decreased mRNA expression compared to controls (as observed for *SERPINE1, FN1*, and *ACTA2*). In contrast to *TGF-β*, while its expression levels remained unchanged, the longitudinal analysis revealed that, in control kidneys, expression increased, while in continuously treated slices, it exhibited a mild decrease in every experiment.

Quantification of pro-collagen 1a1 secretion in the perfusate and culture medium revealed a significant increase during NMP in the galunisertib-treated kidney group (p=0.0449) (Figure 4F). This was followed by a significant decrease at 48 hours in both the continued and discontinued treatment subgroups (p=0.0327 and p=0.0372, respectively). No significant differences were observed between the two treatment groups.

### Pharmacokinetics

Finally, we investigated the pharmacokinetics of galunisertib administration during *ex vivo* perfusion. LC-MSMS analysis demonstrated that galunisertib was steadily cleared in the urine, and through urine recirculation, the absolute perfusate concentration remained stable throughout NMP (Figure 5A). The concentration of galunisertib in the post-NMP flush was similar to the perfusate concentation at the end of the NMP, indicating flush-out of the drug (Figure 5B). Despite equal amounts of galunisertib added to the medium at each time point, absolute values of galunisertib were significantly lower at 48h compared to 24h in the continued treatment group (p=0.0346) (Figure 5C).

**Figure 5:**
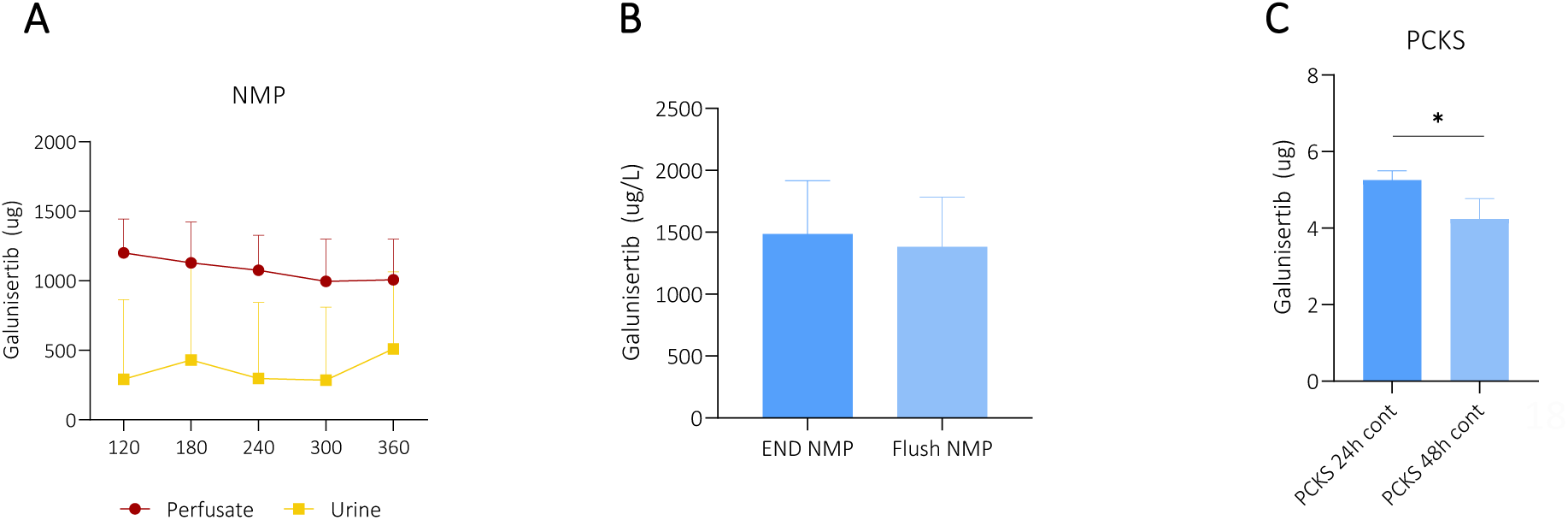
Overview of galunisertib pharmacokinetic study during normothermic machine perfusion, flush, and precision-cut kidney slices A) Longitudinal representation of absolute values of galunisertib in perfusate and urine during NMP (ug); (B) Representation of galunisertib concentration at the end of NMP and during cold flush (µg/L); (C) Representation of absolute values of galunisertib in PCKS medium at 24 and 48 hours of slices with continued treatment (ug). Control slices values were not added due to undetectable results.

## Discussion

One of the biggest hurdles in renal transplantation is the gradual decline in kidney function due to the early onset of interstitial fibrosis and tubular atrophy. ^1,2^ Therefore, identifying and perhaps even inhibiting this phenomenon in a transplantation setting could be of great value. Previous studies have demonstrated that NMP serves as an exceptional platform for targeted and direct drug delivery to organs in multiple animal and human models. ^18–21^ This study explored the potential of delivering galunisertib as an anti-fibrotic agent during *ex vivo* NMP of discarded human kidneys, followed by incubation with PCKS to evaluate long-term effects. It builds on our previous work in a porcine model, where galunisertib was shown to reduce fibrogenesis and inflammation during both NMP and PCKS.^3^

While galunisertib treatment during NMP alone appeared to reduce pro-collagen excretion after 48 hours of tissue incubation, gene expression analysis revealed a more complex response, indicating that a 5-hour exposure during NMP may be insufficient to achieve sustained anti-fibrotic effects. The significant reduction of *TGF-* β, *ACTA2* and *FN1* expression observed after 48 hours with continued galunisertib treatment, but not at earlier time points, indicate that pro-fibrotic human kidneys may require prolonged exposure to galunisertib to obtain anti-fibrotic effects. This contrasts with our previous findings in the porcine model, where 5-hour exposure during NMP alone was sufficient to reduce fibrosis-related mRNA transcripts long-term.^3^ Continued treatment for an additional 48 hours did modulate the expression of fibrosis-associated genes, which aligns with previous PCKS and porcine studies.^3,12,14^ Together, these results support the use of this platform to advance mechanistic insights and guide future therapeutic strategies in human kidney perfusion research.

Importantly, our results demonstrate that galunisertib can be safely administered to human kidneys during NMP without adverse effects on *ex vivo* organ function, viability, or injury. This was supported by consistent functional parameters during NMP, such as renal flow, vascular resistance, urine output, arterial pH, CrCl, and FENa, with no significant differences between the vehicle and galunisertib-treated groups. Furthermore, compound quantification was feasible and revealed stable perfusate concentrations throughout the NMP period. Recent oncological clinical studies report plasma concentrations of galunisertib between 100–1000 ng/mL in patients six hours after a 150 mg oral dose, representing levels approximately 3.7- to 37-fold lower than those used in our study.^23–25^ Despite this higher concentration, no adverse effects were observed, supporting the potential of NMP as a safe and effective platform for *ex vivo* delivery of anti-fibrotic therapies at elevated doses, while minimizing systemic exposure and associated risks.

Our pharmacokinetic analysis indicated that urine recirculation contributed to maintaining stable galunisertib concentrations in the perfusate, as the drug was consistently excreted in the urine. ^7,9,22^ The significant flush-out of the drug after NMP highlights a positive finding as well as raised questions regarding the optimal drug delivery strategy. While the effective removal of galunisertib supports its safety for clinical application by minimizing the risk of systemic exposure in recipients, it also raises the question of whether prolonged drug exposure or omitting the post-perfusion flush might have further enhanced the suppression of fibrogenic gene expression during tissue slice incubation. A similar consideration was raised by Tietjen et al. in 2017,^20^ who concluded that higher poly(lactic acid)–poly(ethylene) glycol nanoparticle concentrations and prolonged exposure times were critical for achieving more effective drug–receptor binding. However, limited literature is available on the time-dependent effects of drug efficacy, particularly in the context of NMP experiments, due to the emerging nature of the field.

Future studies could also explore alternative drug delivery strategies, such as administration during the cold flush or earlier in the preservation process, including HMP, provided that low temperatures do not negatively influence the drug’s solubility and the organ’s enzymatic activity. Another interesting approach could be to explore the use of nanoparticles or other controlled-release mechanisms to guarantee local and long-term effects of the drug in a possible future organ recipient, potentially maximizing the therapeutic benefit and minimizing systemic exposure. Alas, *ex vivo* research still presents a significant gap in translational and transplantation models. ^2^ Lastly, alternative administration routes could be explored, such as via the ureter, intraparenchymal, or subcapsular. Interestingly, during the PCKS experiments, we observed significantly lower concentrations of galunisertib in the culture medium. Given that an equal concentration of the compound was added to the perfusate and medium, this suggests that the drug delivery via diffusion may have resulted in a higher uptake of the tissue. This observation supports the hypothesis that alternative delivery routes could enhance tissue uptake and potentially improve therapeutic efficacy. ^26^

The limitations of this study, such as the small sample size and the lack of comprehensive functional assessments, underscore the need for further investigation. Considerable variability was observed within our experimental groups, particularly in the control group, as reflected in the standard deviations of several key parameters. This variability may have contributed to results not reaching statistical significance. Expanding the sample size and utilizing paired kidneys could help mitigate this limitation and provide a clearer understanding of the therapeutic potential of our intervention. Future studies incorporating larger cohorts, kidney pairs, extended perfusion durations, and refined drug delivery strategies will be essential to validate our findings and advance their translation toward clinical application. In addition, extending the perfusion period could be an avenue worth investigating, as it might allow for more pronounced and sustained therapeutic effects.^2,3,20^ The PCKS model, although useful for monitoring gene expression changes over an extended period, has limitations when it comes to assessing overall kidney function. ^12^

Nevertheless, while limited in its scope, this study provides valuable preliminary evidence suggesting that galunisertib, when administered during NMP, may modulate the expression of key genes involved in fibrogenesis in human kidneys.

The successful translation of our porcine model findings to human tissue opens exciting new avenues for targeted (anti-fibrotic) therapies in the context of organ transplantation, and its different effects between the species highlights the importance of investing in human organ models, as porcine models include young animals without any pre-existing fibrosis. By addressing the limitations identified and further elucidating the mechanisms of action, we can move closer to improving long-term graft survival and expanding the pool of viable donor organs.

## Supporting information

Supplementary Matierial

## Acknowledgements

The authors would like to thank the 34 Lives team, specifically Kathleen St Jean, Stephen Eilert, Paul Wiederhold, Nick Most, and Haley Frost for their hospitality and help performing the experiments. In addition, a big thank the Surgical Research Laboratory staff Susanne Veldhuis, Janneke Wiersma-Buist, and Jacco Zwaagstra for assistance with the biochemical analyses.

