## Supplementary Matierial for "Exploring *ex vivo* modulation of fibrosis in discarded human donor kidneys"

Supplementary Material

### Supplementary Material 1 - Results

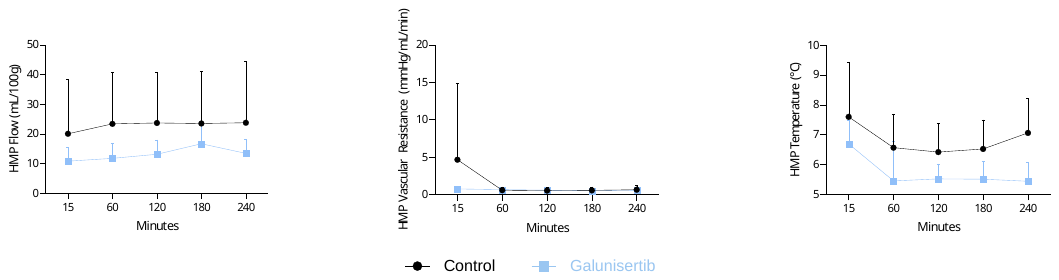

**Supplementary Figure 1:** Overview of functional oxygenated hypothermic machine perfusion (oxHMP) parameters. (A) Grouped longitudinal representation of renal arterial flow (mL/min) per 100g of tissue; (B) Grouped longitudinal representation of vascular resistance (mmHg/mL/min); (C) Grouped longitudinal representation of temperature (˚C).

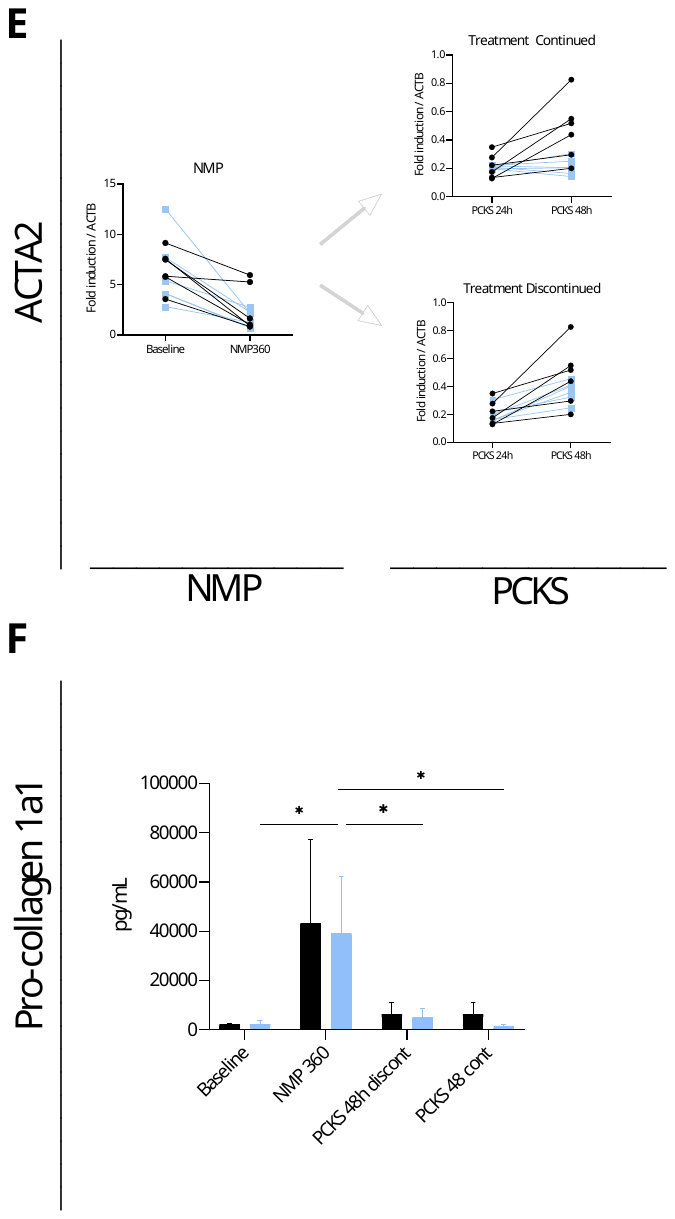

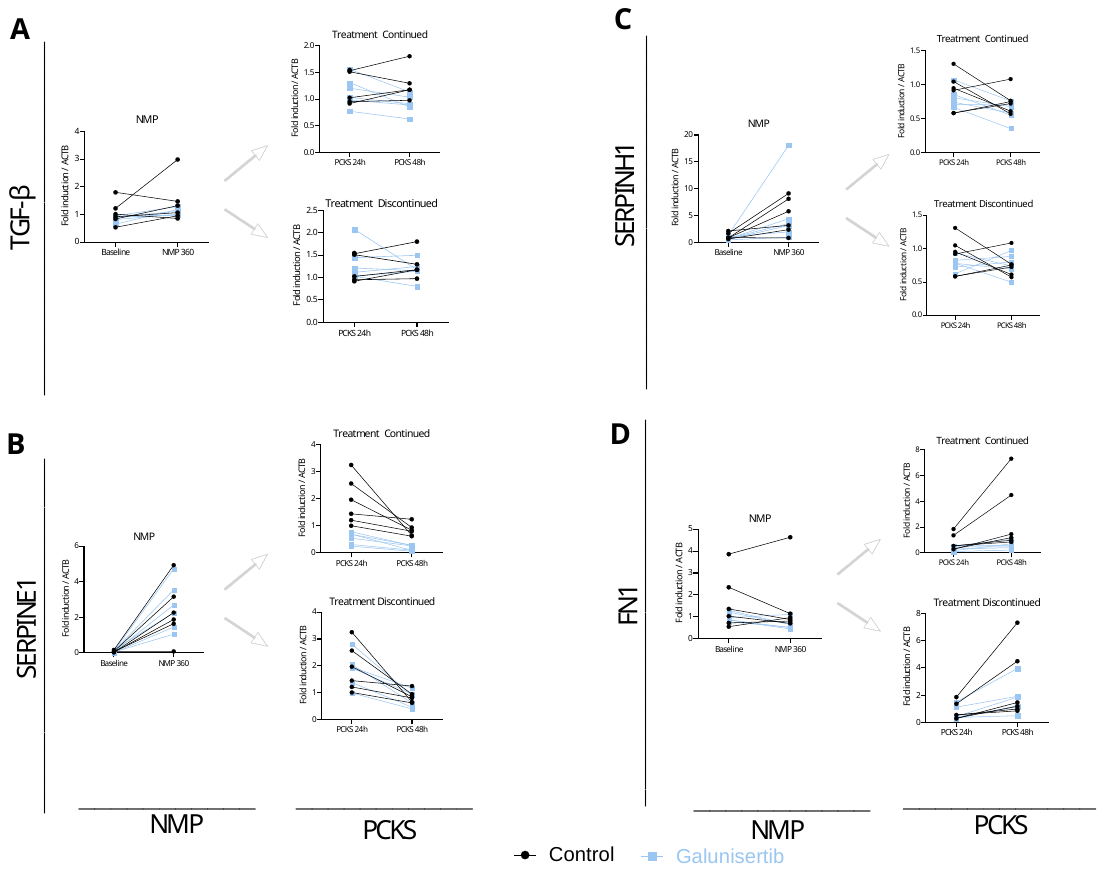

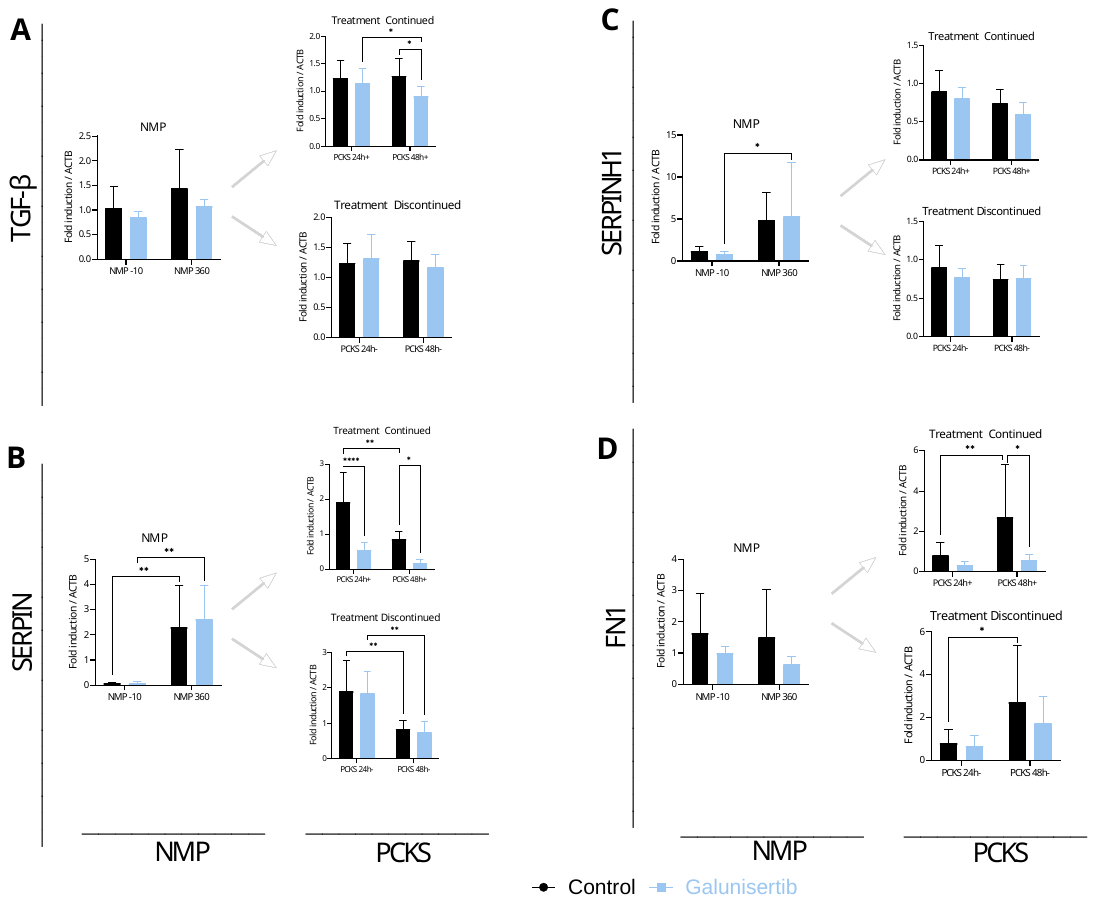

**Supplementary Figure 2:** Overview of fibrosis-related mRNA in tissue and pro-collagen 1a1 protein excretion in perfusate during normothermic machine perfusion and precision-cut kidney slices (continued and discontinued treatment). (A) Representation of *TGF-β* mRNA expression levels in control and galunisertib treated kidneys represented in fold inductions from housekeeping gene *ACTB*; (B) Representation of *SERPINE1* mRNA expression levels in control and galunisertib treated kidneys represented in fold inductions from housekeeping gene *ACTB*; (C) Representation of *SERPINH1* mRNA expression levels in control and galunisertib treated kidneys represented in fold inductions from housekeeping gene *ACTB*; (D) Representation of *FN1* mRNA expression levels in control and galunisertib treated kidneys represented in fold inductions from housekeeping gene *ACTB*; (E) Representation of *ACTA2* mRNA expression levels in control and galunisertib treated kidneys represented in fold inductions from housekeeping gene *ACTB*; (F) Representation of pro-collagen 1a1 protein excretion levels in perfusate during NMP and medium at 48h of incubation from control and galunisertib continued and discontinued treatment kidneys (pg/mL).

### Supplementary Material 2 – Formulas

**Oxygen consumption (VO_2_):**

VO_2_ (mLO_2_/min) = $\left( \left( \left( Hb\times2.4794 \right)+\left( pO2arterial\times K \right) \right)-\left( \left( 0.024794\times Hb\times SO2venous \right)+\left( pO2venous\times K \right) \right) \right)\times Q$

Where Hb is the hemoglobin concentration in mmol/L, pO_2_ is the partial oxygen pressure in kPa, K is the solubility constant of oxygen in water at 37°C and equals 0.0225 (mL O_2_ per kPa), SO_2_ is the saturation in %, Q is the renal perfusate flow (dL/min), and RW is the renal weight in grams.

**Creatinine Clearance (CrCl):**

CrCl (mL/min/100g)= $\left( \frac{\left( \frac{Urine Cr}{Perfusate Cr} \right)*diuresis}{kidney weight} \right)\times100$

Where Urine and Perfusate creatinine are measured in mmol/L, diuresis is measured in mL/min, and the kidney weight is measured in grams.

**Fractional sodium excretion (FENa):**

FENa (%) = $\frac{urine [Na] \times perfusate [creatinine]}{(urine \left[ creatinine \right] \times perfusate [Na])}\times100$

Where Na is the perfusate and urine sodium and creatinine concentration is in mmol/L.

### Supplementary Material 3 – Sample Analysis

#### Biochemistry assay and NMP parameter calculations

Perfusate and urine samples were stored at -80˚C. Biochemistry analysis of the samples was performed at the local clinical biochemistry lab of the University Medical Hospital Groningen. CrCl and FENa calculations were performed with formulas previously described by our group (formulas also provided in Supplementary Material).^8,11^

#### ATP assay

At the time of collection, biopsies were snap-frozen in liquid nitrogen submerged in Sonification Solution (Ethanol 70% with 2mM EDTA M=372.24 g/mol at a pH of 10.9) and then stored at -80 ˚C until analysis was performed. Renal cortical biopsies and tissue slices (processed in triplicate) were homogenized using a Minibead-beater for two 45-second cycles in ice-cold sonification solution. Following homogenization, samples were centrifuged at 16,000 × g for 5 minutes at 4°C. The resulting supernatants were collected for ATP quantification using a Bioluminescence Kit (Roche Diagnostics, Mannheim, Germany). These supernatants were incubated overnight at 37°C to allow evaporation of the solvent. The remaining pellets were reconstituted, and total protein content was determined using the Pierce BCA Protein Assay Kit (Invitrogen). ATP concentrations were normalized to protein content.

#### Gene Expression assay

At collection, biopsies were snap-frozen in liquid nitrogen and stored at -80 ˚C. Total RNA was extracted from biopsies and pooled tissue slices (three slices per sample) using the GeneJET RNA Purification Kit (Thermo Scientific, K0731). RNA concentration was determined using a NanoDrop 1000 spectrophotometer (NanoDrop Technologies, Wilmington, DE, USA), and RNA integrity was evaluated via electrophoresis. Complementary DNA (cDNA) synthesis was performed using M-MLV Reverse Transcriptase (Invitrogen) at 37°C for 50 minutes. Quantitative real-time PCR (qPCR) was then carried out using gene-specific primers (see Tables 1.1 and 1.2), Taq DNA Polymerase (Invitrogen), and the QuantStudio 7 Flex Real-Time PCR System (Applied Biosystems, Bleiswijk, the Netherlands). The amplification protocol consisted of an initial denaturation step at 95°C for 10 minutes, followed by 40 cycles of 95°C for 15 seconds and 60°C for 1 minute. Threshold cycle (CT) values were normalized to the ACTB housekeeping gene and expressed as fold induction.

| **Table 1.1** | | | | |
| --- | --- | --- | --- | --- |
| **Gene** | **Forward sequence (5′ → 3′)** | **Reverse sequence (5′ → 3′)** | **Reference Sequence (Pubmed)** | **Amplicon size (bp)** |
| *TGF-β* | CAATTCCTGGCGATACCTCAG | GCACAACTCCGGTGACATCAA | NM_000660.3 | 72 |
| *Brand: Sigma (KiCqStart® SYBR® Green Primer)* | | | | |

| **Table 1.2** | | |
| --- | --- | --- |
| **Gene** | **Forward sequence (5′ → 3′)** | **Reverse sequence (5′ → 3′)** |
| *SERPINE1* | ATCCACAGCTGTCATAGTC | CACTTGGCCCATGAAAAG |
| *SERPINH1* | GCTAGAATTCACTCCACTTG | CTAGCTCAATTTGGCTCAC |
| *FN1* | TATGAGCAGGACCAGAAATAC | CCATCTTCATTGTTGTCTCTTC |
| *ACTA2* | AGATCAAGATCATTGCCCC | TTCATGCTATTTCCTGTTTGC |
| *ACTB* | GATCAAGATCATTGCTCCTC | TTGTCAAGAAAGGGTGTAAC |
| *Brand: Sigma (KiCqStart® SYBR® Green Primers)* | | |

**Supplementary Tables 1.1 and 1.2:** Primer sequences. Table 1.1 describes the self-designed TGF-β primer, while Table 1.2 describes the primers orders from Sigma, all KiCqStart® SYBR® Green Primers.

#### TNFα and IL-6 assay

Perfusate samples were analysed with a TNFα and IL-6 DuoSet enzyme-linked immunosorbent assay (ELISA) kits (DY210 and DY206, respectively, from Bio-Techne, Abingdon, UK), according to manufacturer's instructions.

#### Pro-collagen 1a1 assay

Perfusate samples were stored at -80 ˚C after collection, and medium samples were snap frozen in liquid nitrogen and preserved at -80 ˚C. Analysis was performed with the Human Pro-Collagen I alpha 1 ELISA Kit (Abcam, ab10966), according to manufacturer's instructions.

#### Galunisertib quantification assay

To quantify galunisertib, a new high-performance liquid chromatography tandem-MS (LC-MSMS) method was developed, drawing on principles from established LC-MS/MS techniques. Sample preparation included protein precipitation with methanol containing a stable isotope-labeled internal standard (SIL-IS), followed by vortex mixing and centrifugation before analysis. ^27,28^ A TSQ Quantiva Triple Quadrupole mass spectrometer (Thermo Fischer Scientific, USA) with a Vanquish Autosampler, Vanquish Horizon Binary Pump, Vanquish Column Compartment, and Vanquish Charger was used for detection after optimizing the gradient elution program, flow rate, and column temperature for galunisertib separation.

The method was validated according to FDA and EMA guidelines for bioanalytical methods, covering parameters such as linearity, selectivity, carry-over, accuracy, precision, dilution integrity, matrix effect, recovery, and stability. A calibration curve was created using galunisertib standards in plasma, spanning concentrations from the lower limit of quantification to the upper limit. Intra-day and inter-day accuracy and precision were assessed using quality control samples, with acceptance criteria of ±15% for bias and coefficient of variation.
